# Muscope: A miniature on-chip lensless microscope

**DOI:** 10.1101/2021.06.09.447680

**Authors:** Ekta Prajapati, Saurav Kumar, Shishir Kumar

## Abstract

In the past few decades, a significant amount of effort has been put into developing different lensless microscope designs. The existing lensless microscopes are capable of offering high resolution and wide field-of-view using super-resolution and computational techniques. But, the employment of macroscopic illumination system and unscalable opto-mechanical components limit their cost-effectiveness, scalability, mass production and on-chip integration. In this work, we report Muscope, an on-chip microscope, which fixes these issues. It extends a few mm in each dimension and comprises of an off-the-shelf electronic assembly. The futuristic microLED display chip is utilised as the light source. Each microLED on the chip functions as a microscopic light source whose position and brightness can be electronically controlled. To demonstrate Muscope, we imaged human blood smear and microbeads of diameter upto 1 μm. We also provide a proof-of-concept of its suitability with super-resolution and field-of-view enhancement techniques, without additional hardware compulsions.

## I. INTRODUCTION

Optical microscopy is indispensable to life sciences and biomedical applications. For such a fundamental and widely used equipment, conventional light microscopes remain costly, complex, bulky, fragile and largely manual. These limitations adversely impact the delivery of proper health care to a large population living in resource poor or hard to access locations.^1^ Even in other settings, the optical microscopes need trained operators and maintenance and have low throughput. An important, yet generally overlooked, short-coming of the conventional microscopes is their inflexibility towards integration into bigger systems, which is an obstacle for a whole class of new devices, automation and process flows.

To fix these flaws, an impressive array of techniques^2–18^ have been devised to simplify the microscope by removing or severely paring down the lens assembly. The resulting lensless microscopes have much smaller sizes. They are also cost effective, easier to operate and support automation, thanks to their electronic makeup. The major downside of removing the lenses is limited resolution, which has been addressed by computational methods and superresolution techniques.^19–28^ The superresolution techniques fuse multiple low resolution (LR), slightly different images to yield a high resolution (HR) image. Many such implementations obtain impressive resolution and a large field of view from devices which can fit in a palm top.

Yet, these clever demonstrations have unsettling aspects. Extended light sources and mechanical arrangements, which are used to capture the LR images for feeding the superresolution pipeline, restrict the reduction in form-factor, cost and scaling production and make the system hard to automate. In many cases, the assembly or fabrication is cumbersome. Almost all the demonstrations treat the microscopes as standalone units, and not as a part to be integrated in more complex systems, like microfluidics chips or robotic hardware. For example, it is inconvenient to use either the conventional or the palm top microscopes for observing multiple locations simultaneously on microfluidic chips or microwell arrays. It is our belief that focusing on these latter use cases can lead to a plethora of applications.

Muscope, the microscope we present here, resolves these concerns. It is the smallest microscope - extending only a few mm in any direction - capable of high resolution imaging. It has no purely optical or mechanical components and can be assembled like a circuit board. All of its operations can be done over electrical interfaces. In particular, it can be software controlled - locally or remotely - and can be tuned online. Because of its small size, multiple Muscopes can be embedded in a small area, and in theory, their sizes can be tuned according to the desired field-of-view. Muscope uses common off the shelf components, making it low cost and robust.

Muscope achieves these feats by replacing the extended light source with the state-of-the-art microLED displays.^29–31^ microLEDs are micron size LEDs available in various colours. The underlying technology allows high brightness LEDs, so that even a single microLED is adequate for imaging. A microLED display contains a rectangular array of such microscopic LEDs, each of which can be addressed individually. These unique features of microLED displays allow Muscope to attain its tiny form, yet provide high resolution imaging, as we show below. Muscope is lensless, therefore the sensor captures holograms, from which real images are reconstructed.^6,9,32–34^ We also show that Muscope field-of-view is only limited by the extent of the microLED display or the imaging sensors. Microscale movement of light source by electronic means is perhaps the most salient feature of Muscope, which allowed us to implement software controlled superresolution. Our data shows improvement signal to noise ratio (SNR) of HR holograms over the as-captured LR ones. As a test, we image a blood smear and microspheres of different diameters on a glass coverslip. Muscope surpasses the existing designs in terms of simplicity and compactness, leading to scalability in cost, use and automation.

Muscope also presents new possibilities. For example, it can be integrated in microfluidic chips for multi-site observation. It can also be hermetically sealed with ease, expanding its scope to extreme environments. By tuning the size of underlying LEDs and the type of lattice as well as its extent, Muscope can be optimised to various applications. Just like the normal LEDs, microLEDs can be illuminated very rapidly which can be useful in fluorescent or time of flight measurements.

## II. EXPERIMENTAL DETAILS

Figure 1 presents the schematic and constituents of the Muscope. It used off-the-shelf electronic components: an image sensor chip and a microLED array display chip which were kept facing each other. A coverslip on which samples were deposited, was positioned between the sensor and display chip as in Fig. 1(e-f). The distance between the display and the sample was z_1_ and that between the sample and sensor was z_2_. A single board computer (SBC) interacted with the display driver board to illuminate a pattern on the display chip as shown in Fig. 1(a-d). This resulted in the sample hologram on the sensor surface. SBC retrieved the hologram from the sensor as programmed. The display chip and sensor surface were covered with glass coverslips (#1.5) to prevent contamination.

**FIG. 1.**
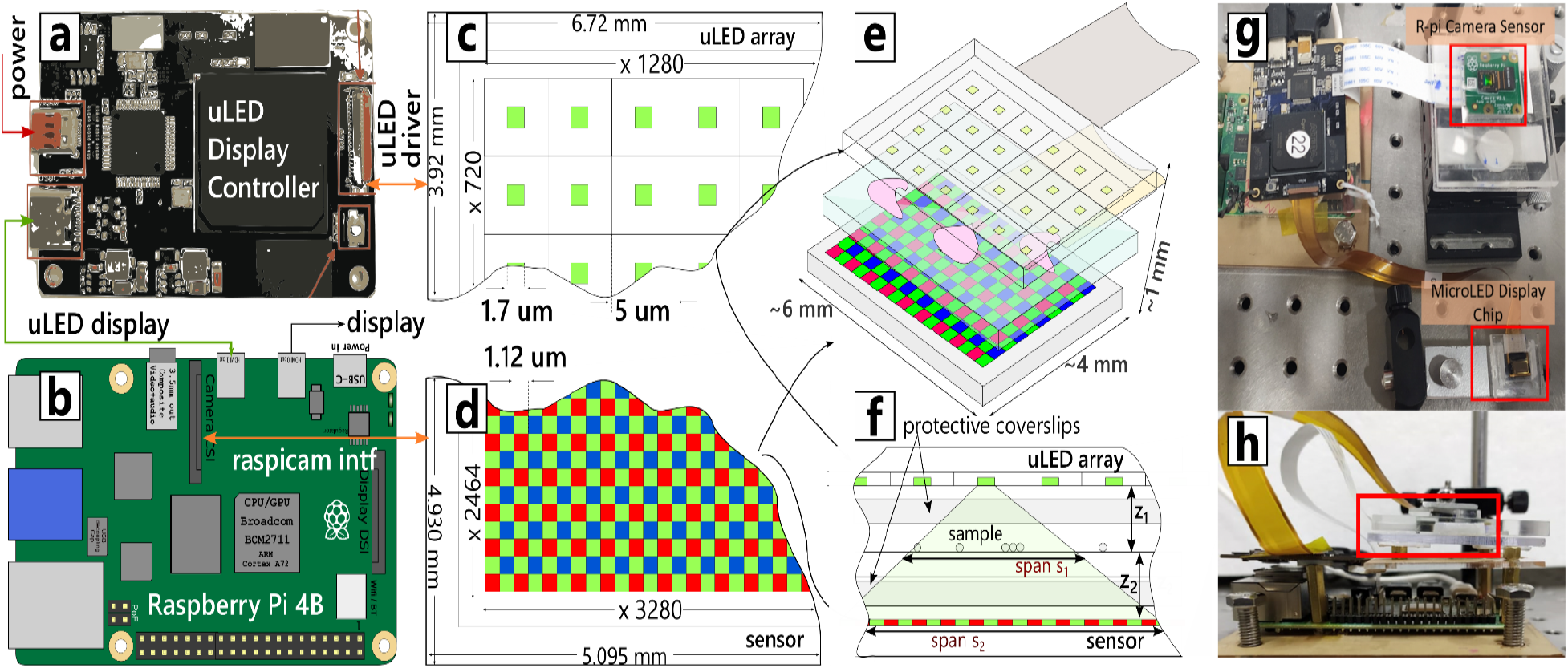
(a-b) microLED display driver and Rpi4 respectively, connected via an HDMI interface. (c-d) Schematic for microLED display and the imaging sensor showing individual cells. (e) Muscope stack, showing the microLED array light source on top and the image sensor at the bottom, sandwiching the sample slide. (f) Part of the sample illuminated by a microLED is magnified at the sensor surface, depending on the distances z_1_ and z_2_. (g-h) Photographs of the setup showing the microLED and the sensor separately and attached together as Muscope.

In our experiments, z_1_ and z_2_ were a few 100 - 1000s of μm. The overall dimensions of Muscope are limited only by the size of display and sensor chips in lateral directions, to about 4 mm × 6 mm in our setup. Along the imaging axis its extent is about 3-4 mm, including the thickness of the sensor and the display.

To move the components with respect to each other for alignment and changing the distances, the display and the sensor were mounted on acrylic enclosures which were affixed to posts on an optical table as in Fig. 1(g-h). The constituents of the Muscope are detailed below:

### 1. MicroLED Display

The display chip (JadeBird Display, Taiwan, Model JBD5UM720PG) is an array of microLEDs arranged on a rectangular lattice of 1280 and 720 points in x and y directions, respectively, depicted in Fig. 1(c). The emitter diameter is 1.7 μm and the pixel pitch in either direction is 5 μm. The microLEDs functioned as a wideband light source emitting at 520-530 nm with full width at half maximum (FWHM) of 30 nm. The intensity of each individually accessible microLED can be varied upto 256 levels. The luminance of an individual microLED is 1×10^6^ - 3×10^6^ Cd/m^2^, which is about 100X higher than that of a common 1mm LED. The overall display area measured 6.72 mm × 3.92 mm. The chip was bonded to a Cu plate which acted as a thermal sink.

The display chip was connected to the driver board through a flat cable. This board regulated the power to the display chip. The maximum power consumed by the display is 1.032 W at 0.369 A current. Individual microLEDs can consume a maximum of 1.6 μA current.

The same board uses micro-HDMI input to interface to a Raspberry-Pi 4B (Rpi4) SBC, running Debian based computer operating system. Rpi4 treated the microLED display interface as an extended 8-bit monochrome display. To illuminate the microLEDs on the array, an image was displayed in this extended area with corresponding pixels set to the desired intensity level. Refresh rates upto 240 Hz are supported by the driver board. In all the experiments, a single microLED at full intensity was used as a light source.

### 2. Imaging Sensor

The imaging sensor was a Sony IMX219 (3280 × 2464 pixels of size 1.12μm × 1.12 μm) chip contained in the Raspberry Pi (Rpi) Cam V2 module, depicted in Fig. 1(d). The lens assembly on top of the imaging sensor was removed by carefully prying it off with a screwdriver, to expose the sensor surface^35^. The surface was cleaned and sealed off with a glass coverslip using an adhesive. The camera module connected to a MIPI interface on the Rpi4 board via a flexible cable as shown in Fig. 1(g).

A system command was used to capture full resolution raw images from the sensor under default settings. The images encoded JPEG as well as raw data, the latter of which was extracted and used for analysis. Since the sensor surface is covered with a Bayer pattern and the LEDs are emitting in green, the blue and red pixels appear dimmed. They were multiplied by constants (2.05 for red and 2.25 for blue) to compensate for loss of intensity. For image capture, the light source and the sensor were activated in a coordinated manner using a Python script.

### 3. Samples

Polystyrene (PS) beads of sizes : 10 ± 0.2 μm, 5 ± 0.1 μm and 1 μm (Sigma Aldrich) were imaged. Beads were dispersed in polydimethylsiloxane (PDMS) following standard PDMS to curing agent ratio (10:1). This mixture was spin coated on a thin glass coverslip using spin coater (Navson Technologies) @500 rpm for 30 seconds. This layer was immediately covered by gently placing a cleaned coverslip on it. The sample was then left undisturbed for overnight at 60°C. Also, prepared microscope glass slide with human blood smear on it, was purchased.

## III. RESULTS AND DISCUSSIONS

Light emitted by a microLED was diffracted by the objects in the sample plane. The diffraction pattern propagates till the imaging sensor, where its intensity variations were captured as hologram by the sensor. Holographic reconstruction algorithms can be used to convert the hologram back to the real image of the sample, even though the phase variations were missing in captured holograms. We imaged several different sized beads (Fig. 2) to ascertain the performance of the Muscope and the reconstruction algorithm.

**FIG. 2.**
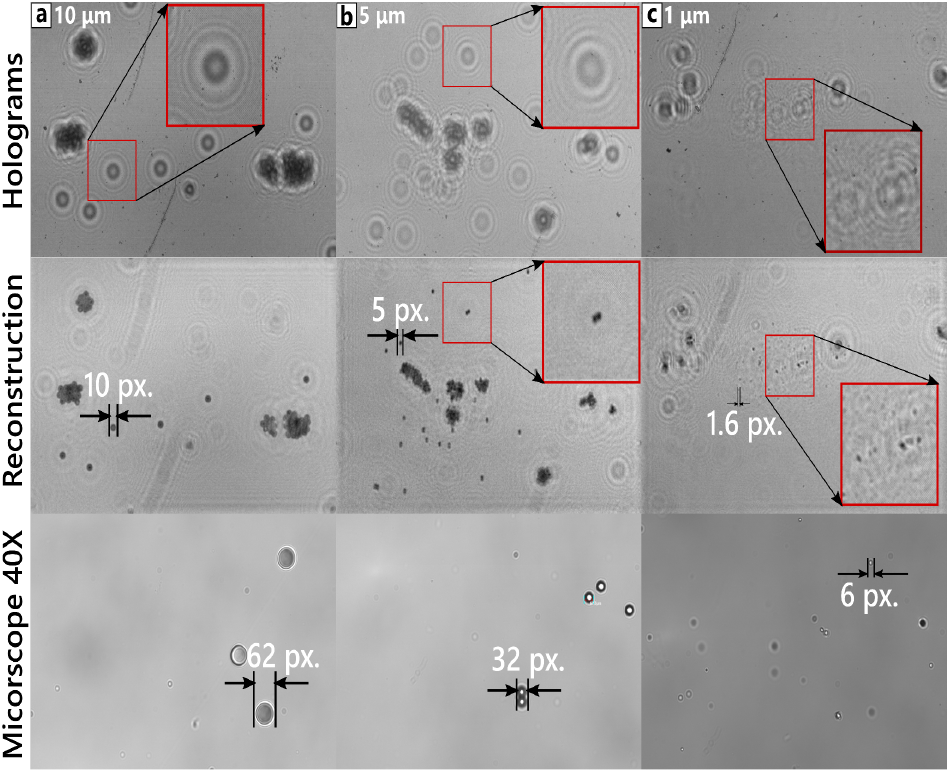
(a-c) Parts of holograms of samples containing beads of specified sizes. The second row shows the reconstruction of real images from these holograms. Bottom row compares to reference images from a 40X microscope objective. Beads in (c) were imaged at z_1_ value smaller than other samples, leading to a higher magnification.

For the imaging setup shown in Fig. 1, if microLEDs were considered as point emitters. Thus, the object plane receives spherical wave (*U*_incident_ = *exp*(*ikr*)*/r*), *r* = (*x, y, z*) and *z* is the imaging axis). Under the assumption that the Fresnel’s number *N*_f_ = *n*(*objectsize*)^2^*/λz*_2_ is small, it can be shown that the hologram captured by the sensor is given by eq.1,^32^

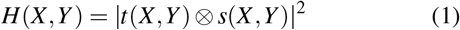

where *R* = (*X,Y,Z*) are coordinates in sensor plane, *t*(*x, y*) is the transmission function of the object. Fresnel’s function *S*(*u, v*) = *exp*(−*iπλz*(*u*^2^ + *v*^2^) acts as a propagator and ⊗ is the convolution operator. The hologram also gets magnified by a factor *M* = (*z*_1_ + *z*_2_)*/z*_1_.

A reconstruction procedure propagates the intensity pattern *H*(*X,Y*) back to the object plane to find the transmission function of the object. The absence of phase information from the hologram makes this process imperfect, in particular, leading to twin images. An interferometric phase retrieval algorithm^36^ deals with these problems and was used here for image reconstruction. The algorithm works well for near field holograms which have Fresnel’s number N_f_ ~ 1 or lower. For Muscope setup, this condition is satisfied readily with objects of size 10 μm or lower. In near field imaging, the intensities are dominated by diffraction terms and the real and virtual images are not well separated out. The iterative algorithm clips one of the twin images in appropriate planes to reduce the effect of the other. The algorithm first generates the virtual image from the intensity of the captured hologram by forward propagating it to a distance z_2_. The object support is detected in this reconstructed image and is replaced by the average of the reconstructed image inside the support. The reconstruction is then back propagated to the object plane (−2z_2_). A gradual clip operator is applied to the generated image to set the field outside the support equal to background. The resulting image is forward propagated to the virtual image plane again (2z_2_). The signal inside the support is again replaced by the average calculated in initial pass. The image is again back propagated to the object plane to get a reconstruction. The process converges after several iterations. For our reconstructions, 5-10 iterations were sufficient. As noted above, the forward and backward propagation used spherical waves under Fresnel-Kirchoff approximation. The value of z_2_ which yielded sharpest images was used for reconstruction.

The refractive index of the medium was a linearly weighted sum of refractive indices of glass (1.33) and air (1.0), where the weights were the fraction of z_2_ covered by each. Two coverslips in sample to sensor path resulted in glass thickness of 340 μm.

In Fig.2, the reconstructed images are compared with the reference images of beads obtained from an inverted microscope with 40X objective. The size of the beads is also shown in pixel units. The sizes of the beads were calculated by manually selecting the areas around the beads of interest and processing them using blob detection and size estimation routines from SciKit skimage libraries.^37^ Average of the size of at least four beads yielded the sizes that are shown.

For the reference images, the physical size of the beads is proportional to the size of the beads in pixels. The same holds true for bead sizes derived from reconstructed images of bigger beads (10 μm and 5 μm, Fig.2(a,b)). The holograms of 1 μm beads (Fig.2(c)) were captured with a smaller z_1_ resulting in a magnified reconstructed image. The magnification factor is derived below, and the resulting size of 1 μm beads approximately follows the size trend of the bigger beads.

Muscope provides another way to calculate the magnification factor using the shift in the holograms when microLEDs at different locations are illuminated. The schematic in Fig.3(a) indicates that the ratio of the shift in the location of the hologram of a bead on the sensor (t_2_) to the shift in location of microLED (t_1_) is the same as the ratio z_2_/z_1_. The frames of the sensor and the microLED display may be oriented at an angle *θ*. We captured a series of holograms of 5 μm and 1 μm beads by illuminating a 13×13 rectangular grid of microLEDs on the microLED display. Closest neighbouring microLEDs in the grid were separated by 2 microLEDs, giving t_1_ = 15 μm. The grid was located in the center of the microLED display, so as to minimize the angle of incidence on the beads. With this arrangement, an algorithm described by Feinup et al.^38^ and coded in Scikit skimage libraries was used to calculate the distances (t_2_) and angles (*θ*) between holograms captured by illuminating neighboring microLEDs. The data is shown in Fig.3(b) for 5 μm and 1 μm beads. The ratio z_2_/z_1_, derived from these plots suggests that M = 1.13 and M = 1.33 for 5 μm and 1 μm beads, respectively. These values are close to those obtained from Fig 2. The shift measurements hold for larger gaps between the neighbouring illuminated microLEDs of 10, 15 and 20 microLEDs - t_2_ scaled almost linearly with the gaps.

**FIG. 3.**
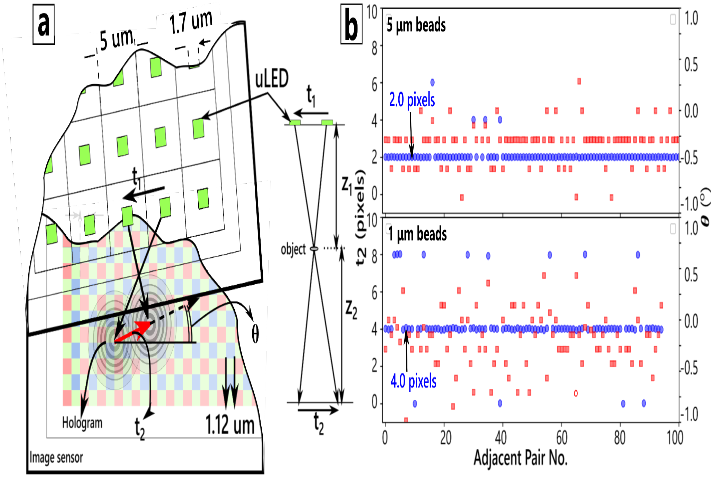
(a) Schematic showing the change in hologram location as the sample is illuminated by adjacent microLED. (b) Plots of the distance t_2_ and angle *θ* for a holograms obtained from pair of adjacent microLEDs. The holograms were obtained from the same setup as in Fig. 2 for 5 μm and 1 μm beads.

The ability to change the location of the light source over large distances over the sensor span is perhaps the most outstanding feature of Muscope. Here we demonstrate two salient advantages of this feature.

First, the holograms captured by illuminating microLEDs separated over large distances enable a field of view only limited by the extent of the microLED display and the sensor. In Fig. 4(a-b), a hologram of the 10 μm bead sample and its zoomed portion at the edges are shown. The illuminating microLED sat on the left hand side of the microLED display. Therefore the beads on the sample located at the farther end on the right, received light at a higher angle, resulting in the hologram getting distorted there (Fig. 4(b)). To image this edge properly a microLED on the right hand side was illuminated (Fig. 4(d)), improving the hologram of the desired area and its reconstruction (Fig. 4(e-f)). An immediate use is to construct large field of view images by fusing non-distorted parts of many holograms obtained from illuminating a series of microLEDs.

**FIG. 4.**
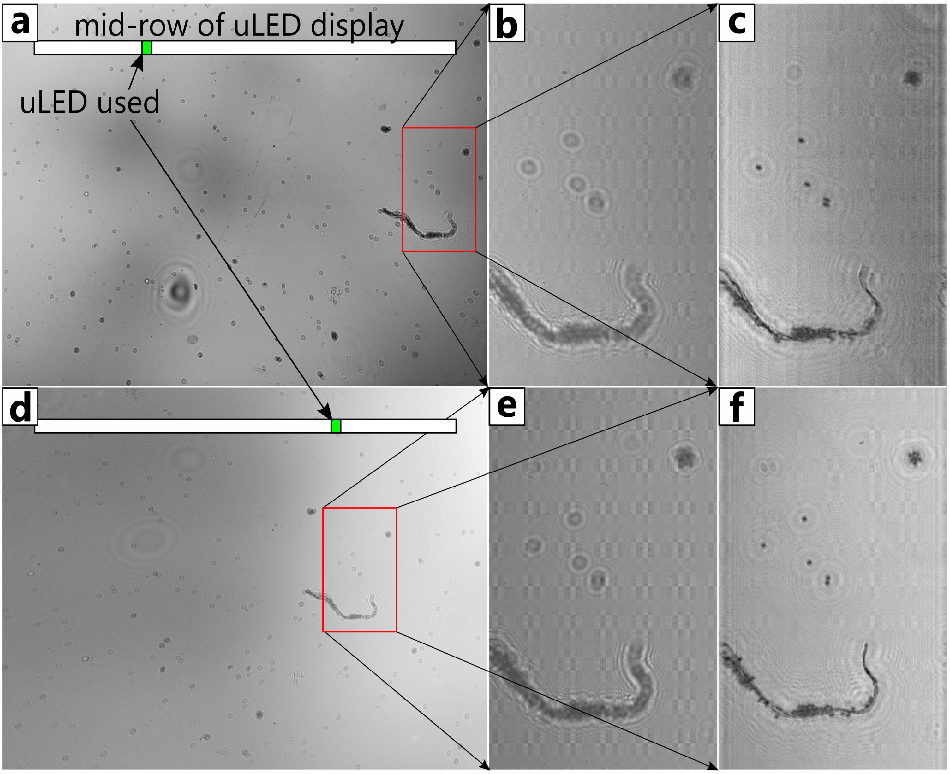
(a) Full frame hologram of 10 μm beads sample using microLED located at the position 335 in a row. (b) At the edge, the hologram shows oblique fringes due to large angle illumination. (c) Reconstruction of the same showing distorted beads. (d) Full frame hologram using microLED at position 923. (e) Hologram of the same area as (b) shows very low distortion, (f) its reconstruction.

Second, the movement of the light source can be used to implement several superresolution techniques available in the literature.^20,39^ The resolution of the reconstructed images depends on several factors. These factors include spatially incoherent and wideband emission from the microLEDs, finite sensor pixel dimensions, low signal to noise ratio (SNR) and approximations made in the reconstruction algorithm. The SNR is affected by the sensitivity of the sensor, the absorbance of the Bayer layer, the illumination intensity from the microLEDs and several noise sources. A combination of these factors limits the number of fringes in the captured hologram of an object. The outermost fringes of the hologram contain the high frequency components and therefore, the finer features of the sample. Even with highly coherent and high intensity lasers, the fringes which are only as fine as a multiple of the sensor pixel size can be analysed (Nyquist criterion).

The impact of finite pixel size of the sensor can be reduced by a multi-image super resolution algorithm, FastSR, described by Elad et al.^20^ It fuses several low resolution but subpixel shifted images to make a high resolution one. With Muscope, the low resolution images were obtained by shifting the position of the microLED on the display array. The change in position of microLED caused a shift in the holograms on the image sensor as shown in Fig. 3(a). With the knowledge of the size of the sensor pixel, the pitch of the microLED display and the distances involved, one can select illumination to result in known (sub-) pixel shifts in holograms. To obtain a N times higher resolution image, the subpixel shifts were binned into bins of size 1/N, in both directions, with a total of N^2^ bins within a pixel. For each pixel in low resolution images there were N^2^ pixels in the high resolution image, contributed by these N^2^ bins. Normally, an illumination pattern will lead to bins having no images or bins having more than one image. In the latter case, median values of all the images in that bin are used, which is a robust estimate for the pixel value. In the former case, an interpolation scheme is needed.

The shift between a pair of holograms from neighbouring illuminated microLEDs was determined by the algorithm of Feinup et al.^38^ The algorithm allows detection of sub-pixel shifts located in bins, whose size can be controlled by passing the upsample factor N to the algorithm.

In the second step of the superresolution algorithm, the effect of the point spread function of the sensor pixels is removed by a gradient descent scheme based on a L_1_ norm. The presence of noise and the need for interpolation is also taken into account in this step. The point spread function of the image sensor has also been determined.^40^ Fig. 5 compares the improvement brought out by the FastSR algorithm compared to a low resolution image. The low resolution images were the same sets of holograms captured from illuminating a 13×13 grid that was described in Fig 3. An upsample factor of 4 was used, the results for higher upsampling were similar. Intensity profile sections across an isolated hologram in low and high resolution are compared in the insets of Fig. 5(a-b). The insets clearly show the increased number of sharply defined fringes in the high resolution hologram. However, only a couple of extra fringes appear in the high resolution hologram, yielding a marginal improvement in resolving two closeby beads in the reconstructed images (insets in Fig. 5(c-d)). Improvement in coherency of the light source, magnification and the SNR is needed to get pixel-limited resolution.

**FIG. 5.**
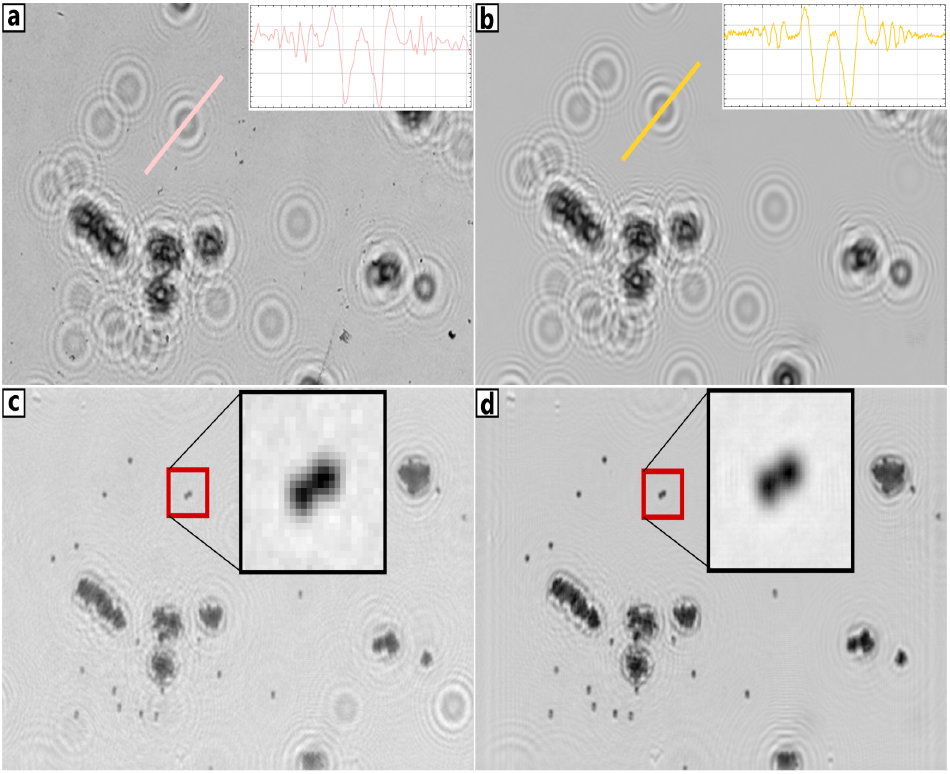
(a) The inset shows the intensity variation across a hologram of a 5 μm bead. (b) The intensity profile has less noise and more fringes for a superresolved hologram. (c) Reconstruction of (a). In-set shows zoomed view of two nearby beads. (d) Reconstruction of superresolved hologram of (b) and showing improved definition of the beads in comparison with (c).

The fusion of images led to the reduction of noise due to averaging of many images and the removal of artefacts which are located at surfaces different from the beads. A couple of dust particles seen on the lower right hand side of the low resolution hologram in Fig 5(a) disappeared in the high resolution image in Fig 5(b). To look at the real world performance of Muscope, we imaged a slide with blood smear on it. A 60X reference image from a microscope, the captured hologram and its reconstruction are displayed in Fig. 6. The reconstructed image is close to the reference image showing the platelets in the same density and sizes as the reference images.

**FIG. 6.**
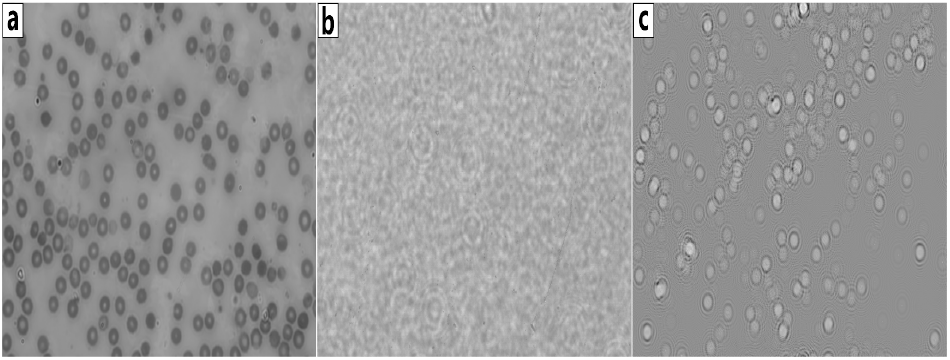
(a) Micorscope image with 60X objective of a blood smear slide. (b-c) Hologram of the same sample and its reconstruction, respectively.

## IV. CONCLUSION

The advancements in electronics and computational techniques have greatly supported lensless imaging systems. However, the light sources used in current lensless microscopes have remained macroscopic and thus, none of the existing implementations is truly “on-chip”. We used a recently developed microLED display chip as the light source to get around this problem. The key aspect of these displays is the ability to change the location and intensity of the individual micron sized emitters over large distances to cover the sample. Using microLED display, we designed and experimentally demonstrated Muscope, an on-chip microscope. Muscope was able to image upto 1 μm diameter beads with just off-the-shelf electronic components, without any optimisations. Changing the locations of the emitters on the display allowed crucial enhancements in resolution and field-of-view, avoiding any optical or mechanical additions to the system. We had also imaged human blood smear to show its aptness with biological samples.

In addition to being the smallest microscope with form factor 3-4mm, Muscope renders multiple future possibilities, especially due to its compatibility with microfluidic technology and robotic technology. Infact, it can be hermetically sealed in a microfluidic device such that the channel lies between the microLED and image sensor chip. The main limitations of Muscope arise from the limited SNR, which is yet to be tackled.

## References

1 Y. Wu and A. Ozcan, “Lensless digital holographic microscopy and its applications in biomedicine and environmental monitoring,” Methods 136, 4–16 (2018).

2 S. B. Kim, H. Bae, K.-i. Koo, M. R. Dokmeci, A. Ozcan, and A. Khademhosseini, “Lens-free imaging for biological applications,” Journal of laboratory automation 17, 43–49 (2012).

3 X. Heng, D. Erickson, L. R. Baugh, Z. Yaqoob, P. W. Sternberg, D. Psaltis, and C. Yang, “Optofluidic microscopy—a method for implementing a high resolution optical microscope on a chip,” Lab on a Chip 6, 1274–1276 (2006).

4 A. Ozcan and U. Demirci, “Ultra wide-field lens-free monitoring of cells on-chip,” Lab on a Chip 8, 98–106 (2008).

5 S. Seo, T.-W. Su, D. K. Tseng, A. Erlinger, and A. Ozcan, “Lensfree holographic imaging for on-chip cytometry and diagnostics,” Lab on a Chip 9, 777–787 (2009).

6 O. Mudanyali, D. Tseng, C. Oh, S. O. Isikman, I. Sencan, W. Bishara, C. Oztoprak, S. Seo, B. Khademhosseini, and A. Ozcan, “Compact, light-weight and cost-effective microscope based on lensless incoherent holography for telemedicine applications,” Lab on a Chip 10, 1417–1428 (2010).

7 S. V. Kesavan, F. Momey, O. Cioni, B. David-Watine, N. Dubrulle, S. Shorte, E. Sulpice, D. Freida, B. Chalmond, J.-M. Dinten, et al., “High-throughput monitoring of major cell functions by means of lensfree video microscopy,” Scientific reports 4, 1–11 (2014).

8 R. Stahl, G. Vanmeerbeeck, G. Lafruit, R. Huys, V. Reumers, A. Lambrechts, C.-K. Liao, C.-C. Hsiao, M. Yashiro, M. Takemoto, et al., “Lens-free digital in-line holographic imaging for wide field-of-view, high-resolution and real-time monitoring of complex microscopic objects,” in Imaging, Manipulation, and Analysis of Biomolecules, Cells, and Tissues XII, Vol. 8947 (International Society for Optics and Photonics, 2014) p. 89471F.

9 S. Amann, M. von Witzleben, and S. Breuer, “3d-printable portable open-source platform for low-cost lensless holographic cellular imaging,” Scientific reports 9, 1–10 (2019).

10 B. Patiño-Jurado, J. F. Botero-Cadavid, and J. Garcia-Sucerquia, “Cone-shaped optical fiber tip for cost-effective digital lensless holographic microscopy,” Applied optics 59, 2969–2975 (2020).

11 E. McLeod and A. Ozcan, “Unconventional methods of imaging: computational microscopy and compact implementations,” Reports on Progress in Physics 79, 076001 (2016).

12 X. Cui, L. M. Lee, X. Heng, W. Zhong, P. W. Sternberg, D. Psaltis, and C. Yang, “Lensless high-resolution on-chip optofluidic microscopes for caenorhabditis elegans and cell imaging,” Proceedings of the National Academy of Sciences 105, 10670–10675 (2008).

13 J. K. Adams, V. Boominathan, B. W. Avants, D. G. Vercosa, F. Ye, R. G. Baraniuk, J. T. Robinson, and A. Veeraraghavan, “Single-frame 3d fluorescence microscopy with ultraminiature lensless flatscope,” Science advances 3, e1701548 (2017).

14 G. Zheng, R. Horstmeyer, and C. Yang, “Wide-field, high-resolution fourier ptychographic microscopy,” Nature photonics 7, 739–745 (2013).

15 M. Rostykus, M. Rossi, and C. Moser, “Compact lensless subpixel resolution large field of view microscope,” Optics letters 43, 1654–1657 (2018).

16 K. Yanny, N. Antipa, W. Liberti, S. Dehaeck, K. Monakhova, F. L. Liu, K. Shen, R. Ng, and L. Waller, “Miniscope3d: optimized single-shot miniature 3d fluorescence microscopy,” Light: Science & Applications 9, 1–13 (2020).

17 W. Luo, A. Greenbaum, Y. Zhang, and A. Ozcan, “Synthetic aperture-based on-chip microscopy,” Light: Science & Applications 4, e261–e261 (2015).

18 G. Zheng, S. A. Lee, Y. Antebi, M. B. Elowitz, and C. Yang, “The epetri dish, an on-chip cell imaging platform based on subpixel perspective sweeping microscopy (spsm),” Proceedings of the National Academy of Sciences 108, 16889–16894 (2011).

19 R. C. Hardie, K. J. Barnard, and E. E. Armstrong, “Joint map registration and high-resolution image estimation using a sequence of undersampled images,” IEEE transactions on Image Processing 6, 1621–1633 (1997).

20 S. Farsiu, M. D. Robinson, M. Elad, and P. Milanfar, “Fast and robust multiframe super resolution,” IEEE transactions on image processing 13, 1327–1344 (2004).

21 W. Bishara, T.-W. Su, A. F. Coskun, and A. Ozcan, “Lensfree on-chip microscopy over a wide field-of-view using pixel super-resolution,” Optics express 18, 11181–11191 (2010).

22 A. M. Maiden, M. J. Humphry, F. Zhang, and J. M. Rodenburg, “Superresolution imaging via ptychography,” JOSA A 28, 604–612 (2011).

23 G. Zheng, C. Shen, S. Jiang, P. Song, and C. Yang, “Concept, implementations and applications of fourier ptychography,” Nature Reviews Physics, 1–17 (2021).

24 G. Zheng, R. Horstmeyer, and C. Yang, “Wide-field, high-resolution fourier ptychographic microscopy,” Nature photonics 7, 739–745 (2013).

25 W. Luo, Y. Zhang, A. Feizi, Z. Göröcs, and A. Ozcan, “Pixel superresolution using wavelength scanning,” Light: Science & Applications 5, e16060–e16060 (2016).

26 Y. Wu and H. Shroff, “Faster, sharper, and deeper: structured illumination microscopy for biological imaging,” Nature methods 15, 1011–1019 (2018).

27 C. Zheng, D. Jin, Y. He, H. Lin, J. Hu, Z. Yaqoob, P. T. So, and R. Zhou, “High spatial and temporal resolution synthetic aperture phase microscopy,” Advanced Photonics 2, 065002 (2020).

28 M. Wang, S. Feng, and J. Wu, “Multilayer pixel super-resolution lensless in-line holographic microscope with random sample movement,” Scientific reports 7, 1–8 (2017).

29 R.-H. Horng, H.-Y. Chien, F.-G. Tarntair, and D.-S. Wuu, “Fabrication and study on red light micro-led displays,” IEEE Journal of the Electron Devices Society 6, 1064–1069 (2018).

30 X. Zhou, P. Tian, C.-W. Sher, J. Wu, H. Liu, R. Liu, and H.-C. Kuo, “Growth, transfer printing and colour conversion techniques towards full-colour micro-led display,” Progress in Quantum Electronics, 100263 (2020).

31 Y. Liu, K. Zhang, B.-R. Hyun, H. S. Kwok, and Z. Liu, “High-brightness ingan/gan micro-leds with secondary peak effect for displays,” IEEE Electron Device Letters 41, 1380–1383 (2020).

32 T. Latychevskaia, “Iterative phase retrieval for digital holography: tutorial,” JOSA A 36, D31–D40 (2019).

33 S. Barkley, T. G. Dimiduk, J. Fung, D. M. Kaz, V. N. Manoharan, R. McGorty, R. W. Perry, and A. Wang, “Holographic microscopy with python and holopy,” Computing in Science & Engineering 22, 72–82 (2019).

34 C. Trujillo, P. Piedrahita-Quintero, and J. Garcia-Sucerquia, “Digital lensless holographic microscopy: numerical simulation and reconstruction with imagej,” Applied Optics 59, 5788–5795 (2020).

35 http://wiki.raspberrytorte.com/index.php?title=Camera_Module_Lens_Modifcation.

36 G. Koren, F. Polack, and D. Joyeux, “Iterative algorithms for twin-image elimination in in-line holography using finite-support constraints,” JOSA A 10, 423–433 (1993).

37 https://scikit-image.org/docs/dev/auto_examples/features_detection/plot_blob.html.

38 M. Guizar-Sicairos, S. T. Thurman, and J. R. Fienup, “Efficient subpixel image registration algorithms,” Optics letters 33, 156–158 (2008).

39 L. Tian, X. Li, K. Ramchandran, and L. Waller, “Multiplexed coded illumination for fourier ptychography with an led array microscope,” Biomedical optics express 5, 2376–2389 (2014).

40 I. Sencan, A. F. Coskun, U. Sikora, and A. Ozcan, “Spectral demultiplexing in holographic and fluorescent on-chip microscopy,” Scientific reports 4, 1–9 (2014).

